## Supplementary figures and images for "The TLK1-MK5 axis regulates motility, invasion, and metastasis of prostate cancer cells"

### sup Figure 1

Supplementary Figure 1

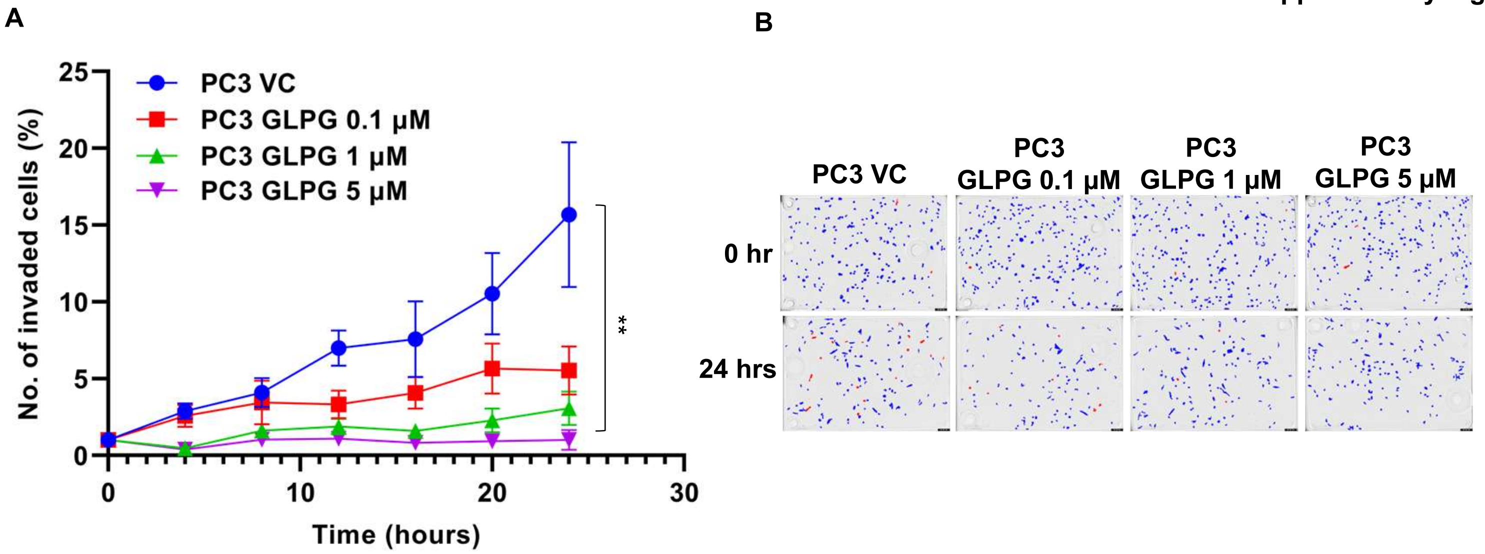
